# A unified spatial transcriptome profiling of ten mouse organs

**DOI:** 10.64898/2026.04.08.715765

**Authors:** Xinyu Ren, Tongxuan Lv, Nanxi Liu, Can Shi, Jinghong Fang, Ning Zhao, Qiang Kang, Dan Wang

## Abstract

Spatial transcriptomics has enabled numerous deep learning models in this area, and training them requires large amounts of high-quality data, especially expression matrices paired with histological images. Here, we present a unified spatial transcriptomic dataset generated using the Stereo-seq platform, covering 10 mouse organs—including brain, kidney, lung, thymus, large intestine, skin, spleen, ovary, testis, and uterus—encompassing 23 tissue sections generated from 21 chips, each with matched ssDNA or H&E staining images. The dataset comprises single-cell-resolution (cell-bin) or square bin-50 (25 µm × 25 µm) expression matrices for each sample, accompanied by corresponding cell type annotations. Annotation robustness was further supported by concordance across different sections of the same tissue and corroboration with canonical marker gene expression patterns. Finally, we compared the characteristics of the cell-bin and bin-50 expression matrices and demonstrated the advantages of cell-bin resolution for cell annotation. This dataset provides a standardized resource for spatial transcriptomics method development, benchmarking, and multimodal analysis.

## Background & Summary

Spatial transcriptomics (ST) has rapidly emerged as a transformative technology, offering critical insights into tissue context and function by combining spatial location information with transcriptome-wide gene expression^1,2^. Among various ST platforms, Stereo-seq stands out due to its subcellular resolution (500 nm) and exceptionally broad transcriptome coverage, as well as its capacity to provide corresponding stained histological images that enable cell segmentation and in-depth study of tissue architecture^3,4^.

Parallel to this, the surge in spatial transcriptomics (ST) technologies has spurred the development of diverse deep learning models and algorithms, particularly large-scale architectures, tailored to decipher these complex multimodal data. They opened new avenues for tasks such as automated cell annotation^5–7^, the decoding of complex cellular interactions^8–10^, and joint analysis of histological images and gene expression matrices^11–13^. Despite these methodological advances, the performance of such models heavily depends on large-scale, high-quality training data. However, the relatively high experimental cost of ST has limited the accumulation of large-scale public datasets—particularly those with unified and standardized profiles^14^. Substantial variability across experimental platforms, laboratory protocols, and data processing strategies has led to pronounced technical heterogeneity, reduced data consistency, and limited cross-dataset comparability. These challenges collectively hinder the development of robust and generalizable deep learning models, highlighting the urgent need for large-scale, unified ST datasets encompassing diverse tissue types^15,16^.

In this study, we present a dataset comprising 10 mouse organs—including brain, kidney, lung, thymus, among others—profiled across 23 sections and 21 chips using Stereo-seq technology (Fig. 1b and Table 1). Each sample is accompanied by corresponding stained histological images (ssDNA or H&E). We systematically assessed image quality to determine whether individual cells could be reliably segmented. For samples with high-quality images enabling accurate cell segmentation, we provide single-cell resolution expression matrices, called “cell-bin”; for the intermediate-resolution profiles, we generated square bins by aggregating spots within a 50 × 50 pixel square region into a single spatial unit (hereafter referred to as “bin-50”) . Using three independent single-cell reference databases (Supplementary Table 1), we performed cell type annotation on both the cell-bin and bin-50 matrices. The annotation results demonstrated consistent spatial distribution within the same tissue and section, as well as concordance with marker gene expression patterns, collectively validating the reliability of the dataset. This dataset provides a valuable resource for deep learning models in spatial transcriptomics. Furthermore, we compared annotation outcomes derived from bin-50 data with those from single-cell resolution data, highlighting the advantages of cell-level expression matrices for refined cell type identification.

**Table 1.** Overview of 10 mouse organ dataset.

| tissue | stain_type | slice | sn_size | kit_type |
| --- | --- | --- | --- | --- |
| brain | ssDNA | slice1 | 1*1 | Stereo_seq_transcriptomics_V1. 3_Kit |
|  |  | slice2 | 1*1 | Stereo_seq_transcriptomics_V1. 3_Kit |
|  |  | slice3 | 1*1 | Stereo_seq_transcriptomics_V1. 3_Kit |
| kidney | H&E | slice1 | 1*1 | Stereo_seq_transcriptomics_V1. 2_Kit |
|  |  | slice2 | 1*1 | Stereo_seq_transcriptomics_V1. 2_Kit |
| lung | ssDNA | slice1 | 1*1 | Stereo_seq_transcriptomics_V1. 3_Kit |
| thymus | H&E | slice1 | 1*1 | Stereo_seq_transcriptomics_V1. 2_Kit |
|  |  | slice2 | 1*1 | Stereo_seq_transcriptomics_V1. 2_Kit |
|  |  | slice3 | 1*1 | Stereo_seq_transcriptomics_V1. 2_Kit |
| large intestine | ssDNA | slice1 | 1*1 | Stereo_seq_transcriptomics_V1. 3_Kit |
|  |  | slice2 | 1*1 | Stereo_seq_transcriptomics_V1. 2_Kit |
| skin | ssDNA | slice1 | 1*1 | Stereo_seq_transcriptomics_V1. 2_Kit |
|  | H&E | slice2 | 1*1 | Stereo_seq_transcriptomics_V1. 3_Kit |
| spleen | ssDNA | slice1 | 1*1 | Stereo_seq_transcriptomics_V1. 3_Kit |
|  | H&E | slice2 | 0. 5*0. 5 | Stereo_seq_transcriptomics_V1. 3_Kit |
| ovary | ssDNA | slice1 | 1*1 | Stereo_seq_transcriptomics_V1. 3_Kit |
|  |  | slice2 | 1*1 | Stereo_seq_transcriptomics_V1. 2_Kit |
| testis | ssDNA | slice1 | 1*1 | Stereo_seq_transcriptomics_V1. 3_Kit |
|  |  | slice2 | 1*1 | Stereo_seq_transcriptomics_V1. 2_Kit |
| uterus | ssDNA | slice1 | 1*1 | Stereo_seq_transcriptomics_V1. 3_Kit |
|  |  | slice2 | 1*1 | Stereo_seq_transcriptomics_V1. 2_Kit |

**Fig. 1.**
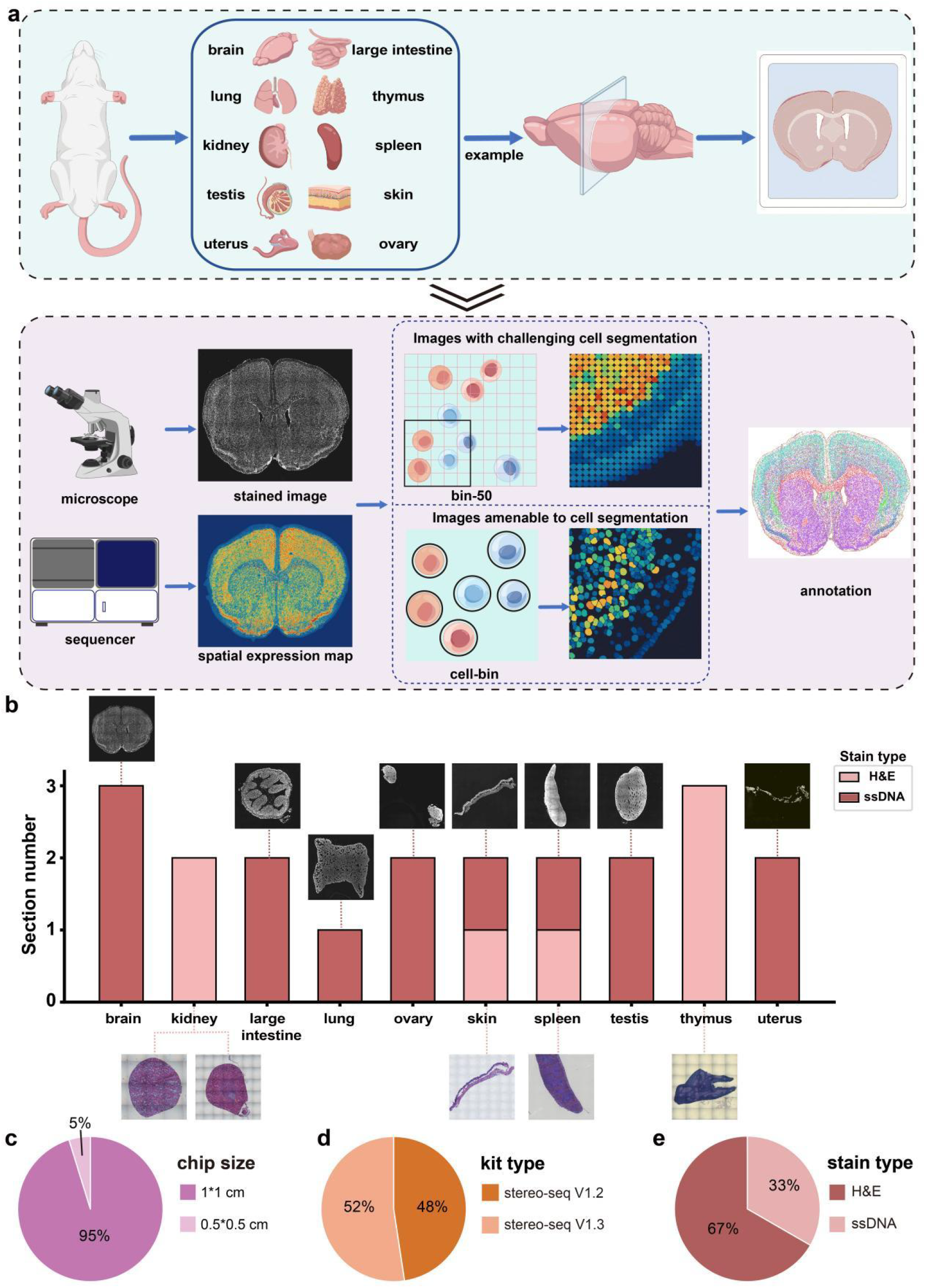
Generation and overview of spatial transcriptomes across 10 mouse organs and 23 tissue sections. (a) Workflow of Stereo-seq procedure and generation of cell-bin and bin-50 expression matrices. (b) Staining types and section numbers for the corresponding tissues. (c) Proportion of different chip sizes used in the dataset. (d) Sequencing kit types used in the dataset. (e) Overview of staining types.

## Methods

### Tissue collection for Stereo-seq experiment

Mouse tissue samples were collected from healthy 12-week-old C57BL/6J mice. All experimental protocols for generating the dataset adhered to ethical regulations regarding animal research and were approved by the Institutional Review Board of BGI (Approval No. BGI-IRB A21001-T1). All methods were carried out in accordance with relevant guidelines and regulations. Fresh-frozen samples were prepared and processed according to the STOmics Stereo-seq Transcriptomics Set User Manual. Briefly, tissues were rapidly dissected, embedded in optimal cutting temperature (OCT) compound, and snap-frozen in liquid nitrogen. Cryosections of 10 μm thickness were prepared using a cryostat and mounted directly onto the surface of Stereo-seq chips, followed by tissue fixation with methanol. This study is reported in accordance with the ARRIVE guidelines (https://arriveguidelines.org).

### Image acquisition and staining

During the imaging stage, an automated microscope equipped with a 10× objective was used for image acquisition. For H&E-stained tissue sections, the epi-brightfield (color camera) mode was selected, whereas for ssDNA-stained sections, the epi-fluorescence mode was used. All images were captured following the imaging requirements specified in the user manual.

### Stereo-seq library preparation and sequencing

Tissue sections placed on the chip were permeabilized with 0.1% pepsin (Sigma, P7000) in 0.01 M HCl at 37 ° C for 10 min. Released mRNA molecules were captured by spatially barcoded probes on the chip surface and reverse transcribed in situ to generate cDNA. The amplified cDNA was then used for DNA nanoball (DNB) generation and library construction. Libraries were sequenced together with coordinate identity (CID) information on the MGI DNBSEQ-Tx platform using paired-end reads.

For samples processed using Stereo-seq V1.2, tissue removal was performed following reverse transcription prior to cDNA release. The released cDNA was subsequently purified and subjected to PCR amplification and purification (details can be found at https://enfile.stomics.tech/STOmics%20Stereo-seq%20Transcriptomics%20Set%20User%20Manual_C%20August%202023.pdf.pdf). For samples processed using Stereo-seq V1.3, the downstream workflow was modified. The separate tissue removal and standalone cDNA purification steps were omitted. Instead, cDNA release and denaturation were carried out using KOH and Elute Additive, followed by neutralization with NS buffer. The resulting cDNA was then directly subjected to partitioned PCR amplification and purification (details can be found at https://enfile.stomics.tech/STUM-TT001%20Stereo-seq%20Transcriptomics%20Set%20V1.3%20for%20Chip-on-a-slide%20User%20Manual.pdf).

### Stereo-seq raw data processing

Stereo-seq V1.2 and V1.3 raw FASTQ data were processed using the SAW (v8.1.0) pipeline^17^. Read 1 contains the coordination identity (CID) and unique molecular identifier (UMI) sequences. CID sequences were aligned to the Stereo-seq capture chip coordinates with up to one mismatch allowed. The corresponding read 2 sequences were mapped to the reference genome (mm10 for mouse) using STAR, retaining only alignments with MAPQ (Mapping Quality) = 255. UMIs sharing the same CID and gene were collapsed into a single count, allowing one mismatch to account for PCR or sequencing errors. Gene expression counts were then aggregated into a spatial profile matrix based on spatial coordinates.

### Cell-bin and bin-50 expression matrices generation

Spatial expression matrices were output as “GEF” files by SAW. For tissue staining images, image processing and cell segmentation were performed using CellBin package^18^, and the resulting CellBin outputs were further subjected to manual inspection and correction. In cases where image quality was insufficient for reliable single-cell segmentation, CellBin was used to align the image with the expression matrix based on track lines. Alternatively, a square bin strategy was adopted to generate a bin-50 expression matrix.

### Downstream processing and cell annotation

The cell-bin and bin-50 “GEF” file were transformed into Anndata format by Stereopy^19,20^. Cell type annotation was performed using the cell2location implementation provided in the STOmics SDAS repository (https://github.com/STOmics/SDAS). Reference single-cell transcriptomic datasets were curated from multiple sources according to tissue type, including the Mouse Cell Atlas^21^ (MCA 3.0; adult mouse), the Tabula Muris Senis^22^ (three-month-old mouse), and the original cell2location publication^23^ (Supplementary Table 1). Cell type annotations were first assessed by comparing cell type distributions across different tissue sections of the same organ to ensure consistency. Resulting annotations were further validated by examining the spatial distribution of canonical marker genes to confirm biological accuracy and spatial coherence.

## Data Records

The raw data have been deposited into the Spatial Transcript Omics DataBase (STOmics DB, https://db.cngb.org/stomics/project) under accession ID STT0000184. All processed data have been uploaded to the CNGB Sequence Archive FTP public service. The dataset includes raw tissue images in .tif format and matrix registered images, raw gene expression matrices in “GEF” format, tissue restricted gene expression matrices in .tissue.gef format, as well as expression matrices at bin-50 or cell-bin resolution, together with the corresponding annotation files. In addition, the raw tissue images in . tif format and the raw gene expression matrices in “GEF” format have been deposited in Zenodo under DOI 10.5281/zenodo.18463527.

## Technical Validation

The spatial transcriptomic dataset presented in this study was generated from freshly dissected mouse organs—including brain, large intestine, lung, thymus, kidney, spleen, testis, skin, ovary, and uterus—comprising a total of 23 tissue sections (Fig. 1a–b). Variations in chip size, kit type, and staining method were present across samples (Fig. 1c, 1d, 1e, and Table 1). Corresponding ssDNA and H&E-stained images were acquired for each tissue section. For samples exhibiting clear, single-cell-resolvable morphology, single-cell expression matrices were generated; all remaining samples were processed to produce bin-50 expression matrices.

The high quality of the resulting expression matrices was validated by their robust transcriptional profiles. At the bin-50 resolution, individual spots across all tissue types consistently contained more than 700 genes and exhibited an average UMI count greater than 1000. Furthermore, at the bin-200 (100 µm × 100 µm) resolution, each spot encompassed over 4500 distinct genes with an average UMI count exceeding 16,000 (Table 2 and Supplementary Table 2). On the other hand, over 76% of the tissue sections exhibited sufficient image clarity for accurate cell segmentation, enabling the generation of cell-bin matrices (Fig. 2a). For these cell-bin expression matrices, the resulting matrices demonstrated well-defined transcriptional profiles, with a median of over 120 genes detected per single cell and an average UMI count exceeding 170 per cell (Table 3).

**Table 2.** Statistics of genes captured at different squarebin sizes. Squarebin Size indicates the number of DNBs combined into a single analysis unit. Mean Gene Types and Median Gene Types represent the average and median number of gene types captured per bin, respectively. Mean UMI Counts and Median UMI Counts denote the average and median unique molecular identifier (UMI) counts per bin, reflecting the levels of gene expression. The table summarizes the statistics for slice 1 of each tissue.

| Tissue | Squarebin Size | Mean Gene Types | Median Gene Types | Mean UMI Counts | Median UMI Counts |
| --- | --- | --- | --- | --- | --- |
| brain | 20 | 137 | 132 | 203 | 183 |
|  | 50 | 716 | 734 | 1,264 | 1,240 |
|  | 200 | 5,070 | 5,537 | 19,599 | 20,619 |
| kidney | 20 | 154 | 147 | 219 | 205 |
|  | 50 | 760 | 751 | 1,360 | 1,282 |
|  | 200 | 4,997 | 5,257 | 20,626 | 18,828 |
| lung | 20 | 498 | 476 | 992 | 922 |
|  | 50 | 2,258 | 2,273 | 6,103 | 5,852 |
|  | 200 | 9,921 | 10,991 | 90,498 | 94,652 |
| thymus | 20 | 152 | 156 | 192 | 197 |
|  | 50 | 775 | 829 | 1,187 | 1,233 |
|  | 200 | 4,878 | 5,778 | 17,745 | 19,101 |
| large intestine | 20 | 749 | 791 | 1,513 | 1,519 |
|  | 50 | 2,812 | 3,122 | 9,227 | 9,203 |
|  | 200 | 8,983 | 10,690 | 130,338 | 132,681 |
| skin | 20 | 145 | 122 | 216 | 166 |
|  | 50 | 703 | 621 | 1,315 | 1035 |
|  | 200 | 4,840 | 5,122 | 18,588 | 16,268 |
| spleen | 20 | 402 | 327 | 1,007 | 778 |
|  | 50 | 1,766 | 1,541 | 6,209 | 4,828 |
|  | 200 | 8,297 | 8,665 | 92,982 | 77,435 |
| ovary | 20 | 668 | 598 | 1,232 | 997 |
|  | 50 | 2,685 | 2,697 | 7,505 | 6,162 |
|  | 200 | 9,336 | 11,038 | 106,801 | 83,634 |
| testis | 20 | 540 | 575 | 740 | 794 |
|  | 50 | 2,284 | 2,475 | 4,577 | 5,003 |
|  | 200 | 9,488 | 10,445 | 69,357 | 77,974 |
| uterus | 20 | 248 | 126 | 442 | 219 |
|  | 50 | 1,103 | 624 | 2,655 | 1,310 |
|  | 200 | 5,610 | 4,729 | 35,443 | 17,155 |

**Table 3.** Statistics of cell bin properties across mouse organs. Cell Count refers to the total number of cells captured in the cell-bin expression matrices. Mean Cell Area and Median Cell Area represent the average and median cell areas, respectively, measured in square micrometers (µm^2^). Mean Gene Types and Median Gene Types indicate the average and median number of unique genes detected per cell. Mean UMI Counts and Median UMI Counts denote the average and median Unique Molecular Identifier (UMI) counts per cell, reflecting the overall transcript abundance.

| Tissue | Slice | Cell Count | Mean Cell Area | Median Cell Area | Mean Gene Types | Median Gene Types | Mean UMI Counts | Median UMI Counts |
| --- | --- | --- | --- | --- | --- | --- | --- | --- |
| brain | slice1 | 113,170 | 185.25 | 171.25 | 290 | 248 | 476 | 385 |
|  | slice2 | 106,238 | 177.5 | 163.5 | 281 | 236 | 504 | 403 |
|  | slice3 | 108,434 | 169.5 | 154.25 | 295 | 248 | 505 | 403 |
| kidney | slice1 | 156,908 | 76.5 | 66.25 | 125 | 95 | 178 | 125 |
|  | slice2 | 66,975 | 76.25 | 68.25 | 129 | 115 | 175 | 151 |
| lung | slice1 | 202,860 | 84.75 | 75.25 | 497 | 446 | 972 | 826 |
| large intestine | slice1 | 28,685 | 92.5 | 80.5 | 860 | 815 | 1786 | 1571 |
|  | slice2 | 29,310 | 86.25 | 75.5 | 283 | 261 | 529 | 466 |
| skin | slice1 | 24,885 | 152.25 | 134 | 249 | 179 | 406 | 258 |
|  | slice2 | 23,805 | 119.5 | 100.5 | 246 | 174 | 424 | 263 |
| ovary | slice1 | 51,698 | 76.75 | 65.25 | 694 | 610 | 1277 | 307 |
|  | slice2 | 46,599 | 76.75 | 65.25 | 209 | 186 | 331 | 307 |
| testis | slice1 | 148,070 | 74.5 | 66.5 | 450 | 421 | 605 | 298 |
|  | slice2 | 140,228 | 83.25 | 72 | 273 | 243 | 382 | 333 |
| uterus | slice1 | 12,941 | 102.5 | 85.75 | 647 | 584 | 1208 | 1039 |
|  | slice2 | 13,325 | 109.5 | 87.75 | 329 | 281 | 855 | 659 |

**Fig. 2.**
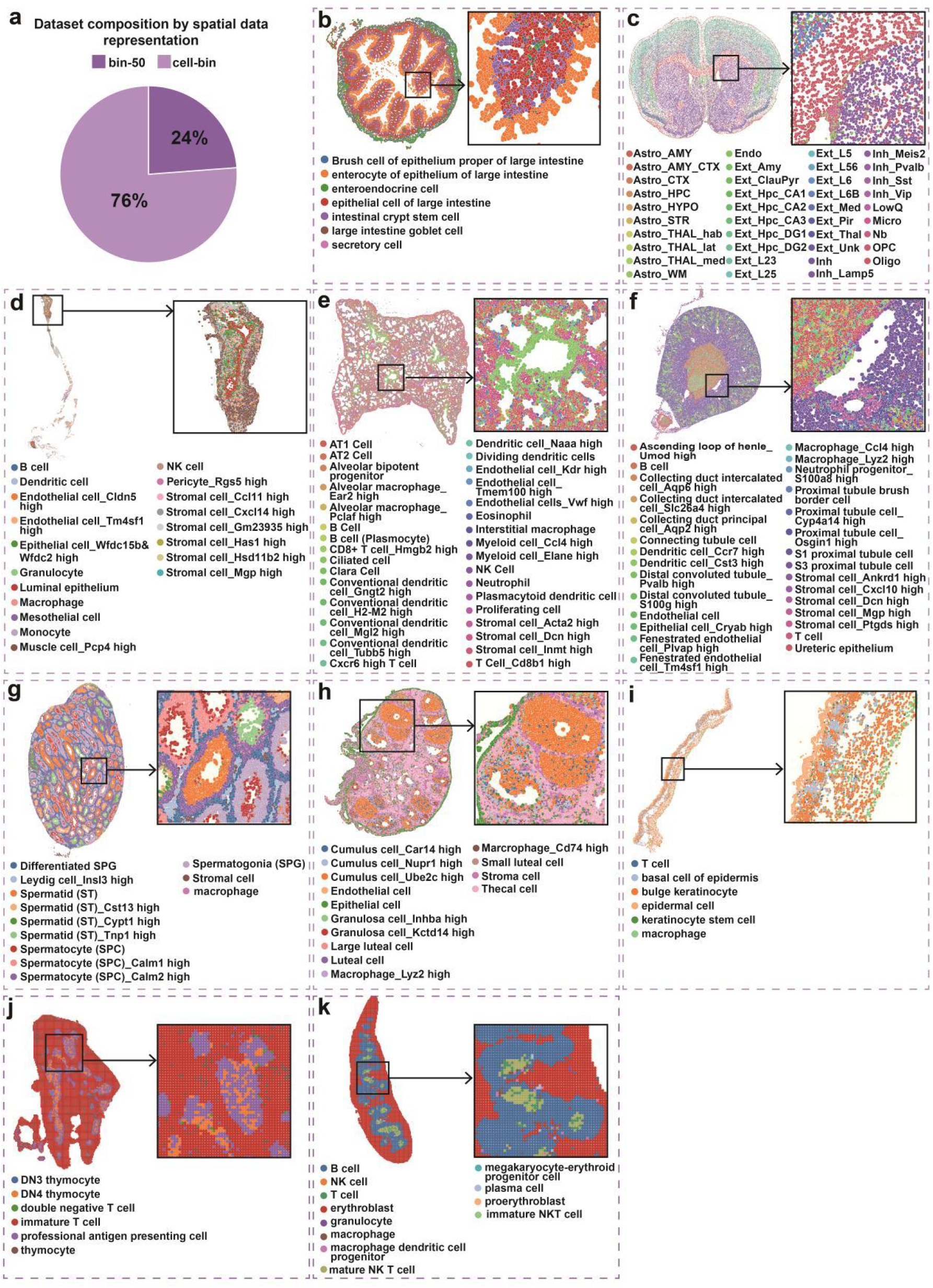
Representative spatial cell type annotations across 10 mouse organs. (a) The proportion of tissue sections analyzed using bin-50 or cell-bin representations. (b–i) Spatial distribution of cell type annotations at cell-bin for representative tissue slices: (b) large intestine slice 2, (c) brain slice 2, (d) uterus slice 2, (e) lung slice 1, (f) kidney slice 1, (g) testis slice 1, (h) ovary slice 2, (i) skin slice 1. (j–k) Spatial distribution of cell type annotations using bin-50 for (j) thymus slice 3 and (k) spleen slice 1.

After removing low-quality spots with low sensitivity, we performed cell annotation on those tissue sections. The resulting cellular annotations accurately delineated the histological architecture, with well-demarcated structural layers consistent with the organ’s anatomy (Fig. 2b–k and Supplementary Fig. 1). For example, in the mouse brain sections, our Cell2location-based annotation clearly resolved distinct neuroanatomical structures, such as the anterior commissure, olfactory limb (aco) and the genu of corpus callosum (ccg) (Fig. 2c and Supplementary Fig. 1e). These spatial patterns are highly concordant with the Allen Mouse Brain Atlas and align with the refined regional annotations reported by Yao et al^24^ (2023). Specifically, our results successfully captured the high-density enrichment of mature oligodendrocytes within these white matter fiber tracts, consistent with the high-resolution MERFISH data from the aforementioned study. Furthermore, in the mouse testis sections, our annotation successfully reconstructed the centripetal distribution of germ cells: spermatogonia (SPG) were precisely localized at the basement membrane, while spermatids were significantly enriched in the center of the seminiferous tubules (Fig. 2g and Supplementary Fig. 1f). Additionally, the mapping results showed that Leydig cells distributed as clusters precisely within the interstitial spaces. These spatial features are highly consistent with the testicular spatial atlas reported by Chen et al^25^ (2021).

As a validation of data robustness, we first assessed the concordance of cell annotations between different slices of the same tissue, and the high cross-slice consistency confirmed the reliability of our cellular annotations as well as the accuracy of both the cell-bin and bin-50 expression matrices across 10 mouse organs (Fig. 2b–l and Supplementary Fig. 1). We then compared the proportional distribution of annotated cell counts and found that cell type compositions were highly consistent among biological replicates of the same organ (Fig. 3a, d). Notably, separate visualization of the most abundant cell type in each organ revealed a consistent spatial distribution of these cells across distinct slices from the same tissue (Fig. 3b, e). To further validate this observation, we examined the expression patterns of canonical markers for each dominant cell type and observed that the marker signals co-localized precisely with the annotated cell positions (Fig. 3c, f, i, l). Specifically, we confirmed the following marker–cell type pairs: Meis2 for Int_Meis2 cells in the mouse brain, Rag1 for immature T cells in the mouse thymus^26^, Spp2 for S1 proximal tubule cells in the mouse kidney^27^, and Slc4a1 for erythroblast cells in the mouse spleen^28^. Importantly, despite morphological variations among different sections of the same tissue, the spatial distribution of the most abundant cell type and its marker genes remained consistent (Fig. 3g–l).

**Fig. 3.**
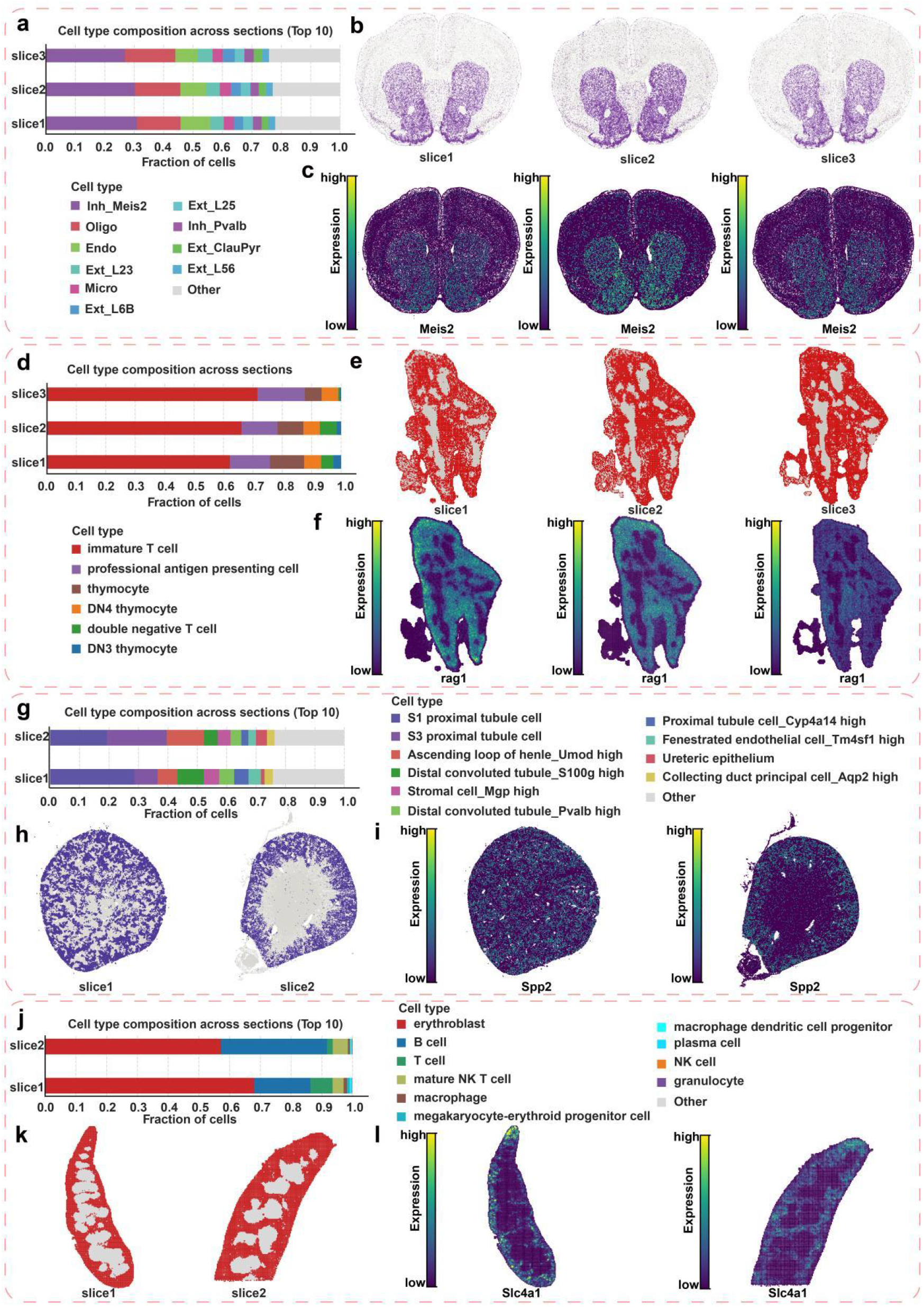
Validation of cell type annotation across representative organs. (a–c) Brain sections (ssDNA, cell-bin): (a) composition of the top 10 annotated cell types, (b) spatial distribution of Inh_Meis2, and (c) spatial expression of its marker Meis2. (d–f) Thymus sections (H&E, bin-50): (d) composition of annotated cell types, (e) spatial distribution of immature T cell, and (f) expression of its marker Rag1. (g–i) Kidney sections (H&E, cell-bin): (g) composition of the top 10 annotated cell types, (h) spatial distribution of S1 proximal tubule cell, and (i) expression of its marker Spp2. (j–l) Spleen sections (slice 1: ssDNA; slice 2: H&E; bin-50) : (j) composition of the top 10 annotated cell types, (k) spatial distribution of erythroblast, and (l) expression of its marker Slc4a1.

We additionally compared the performance of the cell-bin and bin-50 matrices, motivated by their distinct characteristics: the cell-bin matrix provides superior single-cell UMI assignment accuracy, whereas the bin-50 matrix yields higher UMI counts per spot (Table 2 and Table 3). The spatial distribution of cells, as visualized by histological staining, was consistent with that reflected in the cell-bin expression matrix (Fig. 4a–b, Fig. 4e–f, and Fig. 4i–j). Furthermore, we observed a high degree of concordance between the cell-bin and bin-50 annotation results at the global level (Fig. 4c–d, Fig. 4g–h, and Fig. 4k–l). Notably, at the local architectural level, the cell-bin matrix preserved intact hollow structures within the tissue and enabled the identification of additional cell populations, uncovering richer molecular characteristics (Fig. 4c, 4g, and 4k). In the mouse testis, the cell-bin expression matrix successfully annotated macrophages, whereas the bin-50 strategy failed to identify this population within the same localized region (Fig. 4m). Similarly, in the mouse ovary, the cell-bin matrix enabled the precise annotation of endothelial cells and epithelial cells, while the bin-50 strategy failed to capture these signals (Fig. 4n). In localized areas of the mouse kidney, compared to the bin-50 matrix which only identified Cryab high epithelial cells and ureteric epithelium, the cell-bin matrix demonstrated finer resolving power, capturing a more diverse range of cell subtypes within these regions (Fig. 4o).

**Fig. 4.**
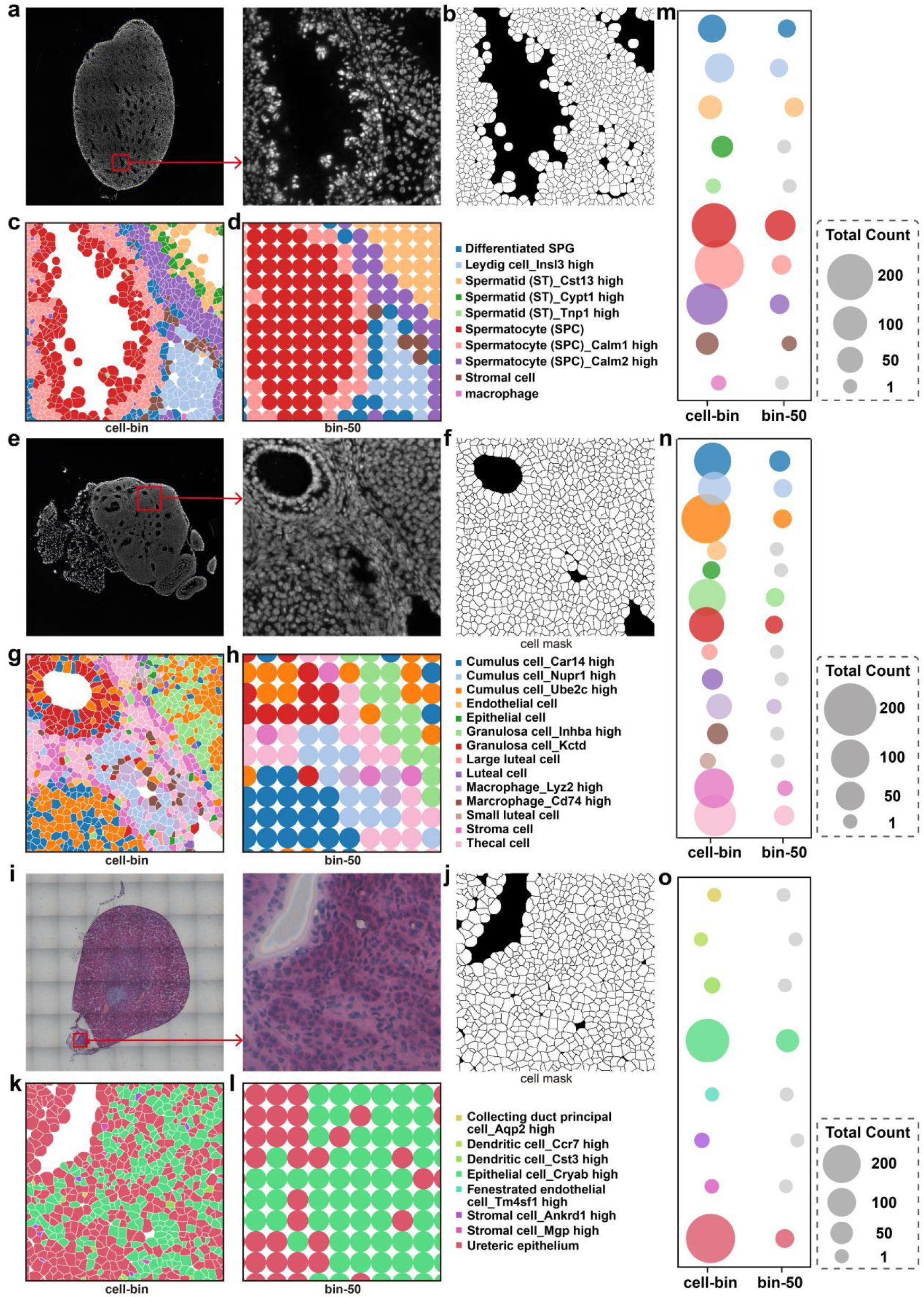
Comparison of spatial expression representation strategies for cell type annotation. (a–l) Cell segmentation and annotation across diverse tissue slices, including (a–d) testis slice 1, (e–h) ovary slice 1, and (i–l) kidney slice 1. For each tissue, panels display the full tissue image with a magnified view of a selected region of interest (ROI) (a, e, i), the corresponding cell segmentation mask (b, f, j), and cell type annotations within the ROI generated using the CellBin (c, g, k) and bin-50 (d, h, l) strategies. (m–o) Quantitative comparison of annotation performance. Bubble plots summarize the cell counts for each cell type within the corresponding magnified ROIs for (m) testis, (n) ovary, and (o) kidney. Bubble size represents the total count of annotated units identified by each strategy. Light grey bubbles indicate cell types not captured under the bin-50–based annotation.

## Code availability

All data preprocessing and analysis workflows were performed using CellBin v2 (https://github.com/STOmics/cellbin2), Scanpy (v1.9.8, https://github.com/scverse/scanpy), and SDAS (https://github.com/STOmics/SDAS).

## Acknowledgements

We thank the China National GeneBank for providing data storage support for this study. This work was supported by the Zhuhai Basic and Applied Basic Research Foundation (2220004002717), the National Natural Science Foundation of China (32400456), and in part by the Guangdong Higher Education Upgrading Plan (UICR0400007-24) at Beijing Normal-Hong Kong Baptist University, Zhuhai, China. We also thank Mei Li, Ying Zhang, Yumei Li, Jing Guo, Zhenzhen Zhu, Yasheng Liu, Dan Zhang for their assistance.

## Author contributions

X.R., Q.K., and D.W. designed the research. X.R., T.L., N.L., C.S., J.F., Q.K., and D.W. have obtained and analyzed the data. X.R., and T.L., drafted the manuscript. X.R., T.L., N.L., N.Z., N.Z., Q.K., and D.W. revised and edited the manuscript. X.R., Q.K., and D.W. supervised the study. All authors have made a substantial contribution to the manuscript.

