## Supplement for "A unified spatial transcriptome profiling of ten mouse organs"

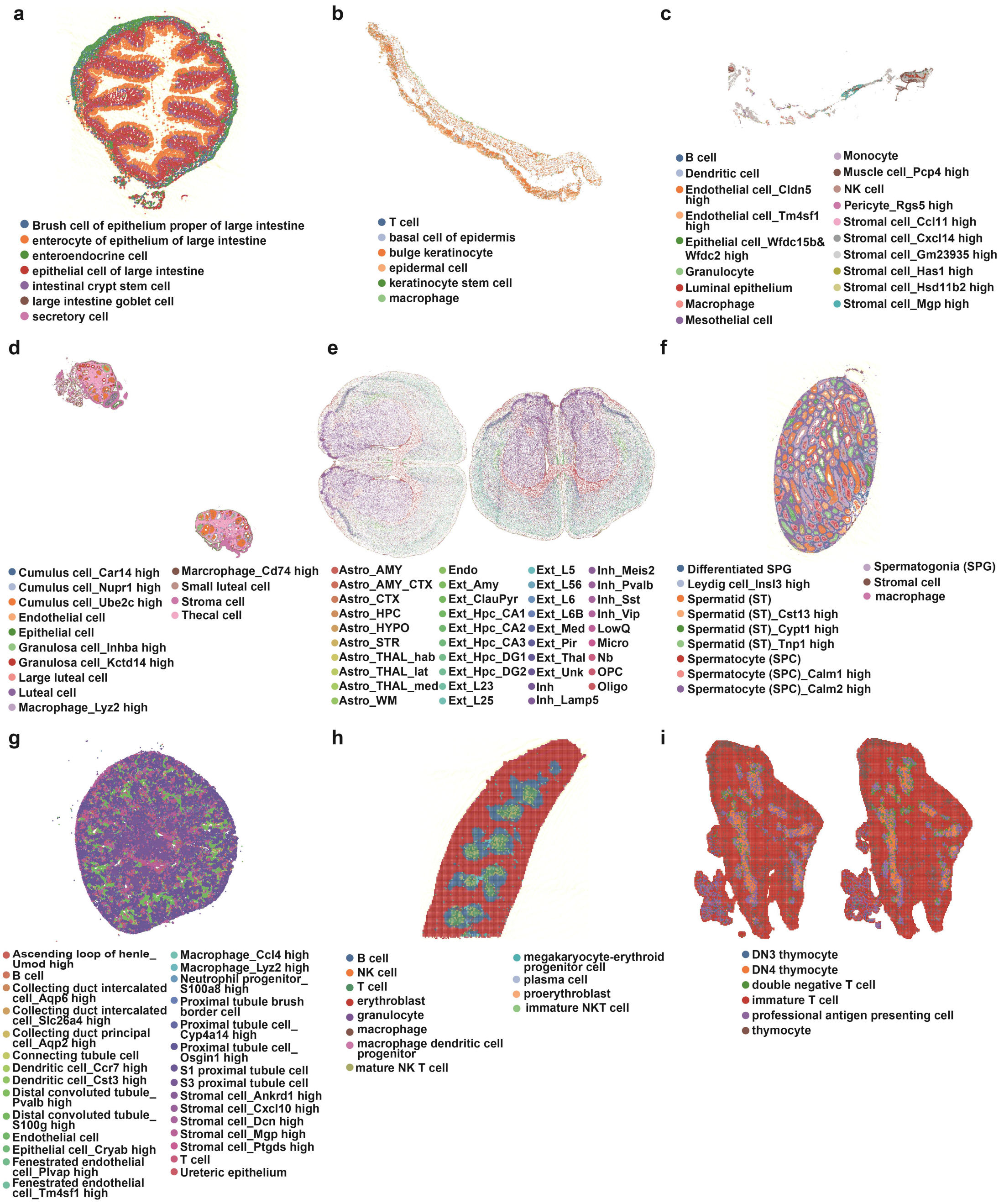


**Supplementary Fig.1 Spatial distribution of cell type annotations across various mouse tissue sections.** (a–g) Spatial distribution of cell type annotations at cell-bin resolution for representative tissue slices: (a) large intestine slice 1, (b) skin slice 2, (c) uterus slice 1, (d) ovary slice 1, (e) brain slice 1 (left) and brain slice 3 (right), (f) testis slice 2, (g) kidney slice 2. (h–i) Spatial distribution of cell type annotations at bin-50 resolution for (h) spleen slice 2, and (i) thymus slice 1 (left) and thymus slice 2 (right).

| tissue | cell-bin/bin-50 | reference |
|---|---|---|
| brain | cell-bin | original cell2location publication |
| kidney | cell-bin | Mouse Cell Atlas |
| lung | cell-bin | Mouse Cell Atlas |
| thymus | bin-50 | Tabula Muris Senis |
| large intestine | cell-bin | Tabula Muris Senis |
| skin | cell-bin | Tabula Muris Senis |
| spleen | bin-50 | Tabula Muris Senis |
| ovary | cell-bin | Mouse Cell Atlas |
| testis | cell-bin | Mouse Cell Atlas |
| uterus | cell-bin | Mouse Cell Atlas |

**Supplementary Table 1. Cell type annotation reference of tissues.**

| Slice | Squarebin Size | Mean Gene Types | Median Gene Types | Mean UMI Counts | Median UMI Counts |
|---|---|---|---|---|---|
| brain-2 | 20 | 138 | 131 | 222 | 198 |
| | 50 | 715 | 726 | 1,383 | 1,353 |
| | 200 | 5,085 | 5,543 | 21,451 | 22,922 |
| brain-3 | 20 | 158 | 155 | 244 | 229 |
| | 50 | 806 | 826 | 1,520 | 1,509 |
| | 200 | 5,397 | 5,815 | 23,595 | 24,365 |
| kidney-2 | 20 | 145 | 158 | 197 | 212 |
| | 50 | 720 | 823 | 1,197 | 1,339 |
| | 200 | 4,393 | 5,557 | 16,461 | 19,868 |
| thymus-2 | 20 | 237 | 245 | 315 | 327 |
| | 50 | 1,161 | 1,258 | 1,944 | 2,029 |
| | 200 | 6,183 | 7,357 | 29,041 | 31,030 |
| thymus-3 | 20 | 198 | 209 | 248 | 258 |
| | 50 | 986 | 1,082 | 1,523 | 1,634 |
| | 200 | 5,609 | 6,490 | 22,493 | 25,688 |
| large intestine-2 | 20 | 299 | 318 | 560 | 591 |
| | 50 | 1,291 | 1,431 | 3,369 | 3,700 |
| | 200 | 6,445 | 7,428 | 46,991 | 53,000 |
| skin-2 | 20 | 145 | 103 | 237 | 153 |
| | 50 | 721 | 564 | 1,445 | 962 |
| | 200 | 4,688 | 4,910 | 20,528 | 14,806 |
| spleen-2 | 20 | 233 | 218 | 311 | 291 |
| | 50 | 1,109 | 1,082 | 1,906 | 1,808 |
| | 200 | 6,013 | 6,771 | 27,632 | 28,174 |
| ovary-2 | 20 | 214 | 215 | 342 | 328 |
| | 50 | 974 | 1,032 | 2,040 | 1,999 |

| | | | | | |
|---|---|---|---|---|---|
| | 200 | 4,915 | 5,721 | 26,044 | 22,322 |
| testis-2 | 20 | 310 | 311 | 437 | 403 |
| | 50 | 1,427 | 1,494 | 2,706 | 2,509 |
| | 200 | 7,379 | 8,049 | 41,134 | 41,106 |
| uterus-2 | 20 | 174 | 127 | 435 | 309 |
| | 50 | 790 | 600 | 2224 | 1867 |
| | 200 | 4645 | 4337 | 34,701 | 24,823 |

**Supplementary Table 2. Statistics of genes captured at different squarebin sizes.** Suffixes -2, and -3 denote slice 2, and slice 3, respectively.
